# Exploiting pathogen defence trade-offs to manage risks of crop pests evolving resistance to biocontrol

**DOI:** 10.1101/2024.10.19.619212

**Authors:** Rosie Mangan, Matthew C. Tinsley, Ester Ferrari, Ricardo A. Polanczyk, Luc F. Bussière

## Abstract

Pathogens often exert strong selection, yet host populations maintain considerable genetic variation for resistance, possibly due to environmental heterogeneity causing fitness trade-offs. These trade-offs could help explain genetic variation for pathogen defence, and also constrain resistance evolution to microbial pesticides (an underappreciated risk). However, the presence and strength of such trade-offs remain unclear. We investigate whether pathogen identity or host diet has a stronger effect on the fitness of resistance alleles. We quantify genetic variation and covariation for pathogen resistance in an insect pest across distinct pathogen and plant diet combinations. We demonstrate substantial heritability, indicating considerable risks of biopesticide resistance. Contrary to conventional thinking in host-pathogen biology, we found no strong genetic trade-offs for resisting two fungal pathogen species, but changes in plant diet dramatically altered selection for resistance, revealing diet-mediated genetic trade-offs affecting pest survival. Our data suggest that trade-offs not strictly related to immunocompetence could nevertheless maintain genetic variation for pathogen defence in natural and agricultural landscapes.

## Introduction

Host populations in natural systems commonly harbour considerable genetic variation for pathogen defence^1–4^. Although classic evolutionary mechanisms like negative frequency-dependent selection (e.g., Red Queen dynamics) can explain some of this variation^5^, genetic variation for infection susceptibility must also be maintained by other evolutionary forces, such as genotype by environment interactions (GEIs)^3,6^. When GEIs exist, a given genotype’s effectiveness in conferring resistance is conditional on specific environmental contexts. This raises a crucial question: which aspects of environmental heterogeneity play the strongest role in driving inconsistent selection on pathogen defence loci^3,7^?

Human activities, particularly in agriculture, often reduce environmental heterogeneity. For example, extensive monocultures diminish landscape variation and increase pest outbreaks^8^, which alongside chemical pesticide use selects for resistant pests and increases crop damage^9,10^. Consequently, there is growing interest in ecologically sustainable insect pest control products, such as microbial biopesticides containing living parasites^11,12^. As the use of these microbial biopesticides increases^13^, so too will selection pressures to evolve resistance against them^14^.

Traditionally, strategies to manage resistance to synthetic pesticides and genetically modified crops involve weakening selection for resistance, for example by minimising pesticide application, employing crop refuges, or rotating use of products with different modes of action^15–17^. An additional approach is Negatively Correlated Cross-Resistance (NCCR), which aims to exploit the fact that resistance to one product sometimes trades-off with resistance to another^18^. However, NCCR has not become an important resistance management tool because resistance to synthetic pesticides often depends on a small number of independent loci, meaning that recombination can rapidly generate genotypes that are resistant to multiple products. Therefore, NCCR has mostly been neglected as a sustainable pesticide resistance-management approach^7,19^.

Here we seek to re-discover the principles behind NCCR and investigate how they can be applied to biopesticides formulated from living microbes; this is especially timely during the ongoing transition to more environmentally sustainable pest control in global agriculture^11^. In contrast to synthetic pesticides, there is much greater potential for NCCR involving biopesticides formulated from living microbes. Defence against the microbes used as biopesticides is typically more genetically complex than for synthetic insecticides or genetically modified crops^20^: indeed in natural systems, pathogen resistance is often highly polygenic and relevant resistance alleles often exist at intermediate frequencies without constant exposure to a particular pathogen^4^. Such complex genetic architecture should make it more difficult for recombination to resolve trade-offs^7^.

Resistance trade-offs may arise due to specific genetic interactions between hosts and parasites: resistance genotypes for one pathogen species or strain can increase susceptibility to another^21^. Leveraging such GEIs could help manage resistance if biopesticides containing different microbial pathogens are used in rotation. In addition to the pathogens themselves, multiple environmental factors, especially variable temperatures, are known to drive GEIs related to pathogen resistance^3,6^. Farmers cannot control temperatures to mitigate resistance risks, yet they do control crop selection, which for polyphagous pests dictates the pests’ diet and can substantially influence infection defence efficacy and immune function^22^. Whether diet can impose heterogeneous selection on resistance genotypes through GEIs remains underexplored. We set out to test the extent to which heterogeneity in both the pathogens used in biopesticides and the crops grown in agricultural landscapes might be used to manage the threats of resistance to microbial biopesticides evolving.

In this study we focus on the cotton bollworm, *Helicoverpa armigera* (Hübner) (Lepidoptera: Noctuidae), a major global agricultural pest with a history of developing resistance to multiple control tactics^23–26^. We study fungal biological control agents because they are especially promising in the context of GEIs: they have a complex infection process involving penetrating the insect cuticle, replicating inside the host, and ultimately killing the host to facilitate onward transmission^27^. The genomes of entomopathogenic fungi encode multiple virulence mechanisms to infect hosts, manipulate their physiology and subvert their defences^28^, while host resistance to these pathogens is typically highly polygenic^20^.

This study has twin aims, one fundamental and one applied. From a fundamental perspective, we asked: to what extent can the pervasive presence of genetic variation for infection defence be explained because GEIs drive inconsistent selection under different environmental conditions? Which is the more powerful driver of inconsistent selection on infection defence genes: variation in the identity of the pathogen, or variation in the diet the animal consumes during infection? We also exploited this evolutionary science for an applied goal. We aimed to quantify the risks that the crop pest *H. armigera* can evolve resistance against fungal biopesticides by quantifying the standing genetic variation for infection susceptibility on which selection could act. Furthermore, we sought to determine how GEIs might be exploited by farmers to make selection for biopesticide resistance inconsistent, thereby managing the threat of resistance evolution to these ecologically sustainable pest control products.

## Results

### Survival consequences of changes in plant diet and pathogen treatment

We used a half-sib breeding design to quantify additive genetic variation for ability to survive pathogen infection; *H. armigera* larvae consumed a diet of leaves from one of three different crop plants and were either uninfected or exposed to one of two different fungal pathogens. Our dataset included 3811 offspring from 37 *H. armigera* sires, collectively mated to 58 dams (1-3 dams per sire). The two pathogen exposure treatments (*Beauveria bassiana* or *Metarhizium anisopliae*) induced markedly higher larval mortality than in uninfected larvae regardless of the leaf diet (Fig. 1). Also, background mortality varied for larvae feeding on the three different food plants: larvae survived best when fed soybean, whereas those reared on tomato or maize were much more likely to die (Fig. 1, Table S1). Interestingly, the ability of fungi to kill larvae was driven by the combination of fungal isolate and larval crop diet (plant:pathogen treatment interaction, parametric bootstrap p-value = 0.003; Table S1): whilst *B. bassiana* caused greater mortality than *M. anisopliae* in larvae feeding on soybean and maize leaves, this virulence advantage disappeared for larvae on tomato (Fig. 1).

**Fig 1.**
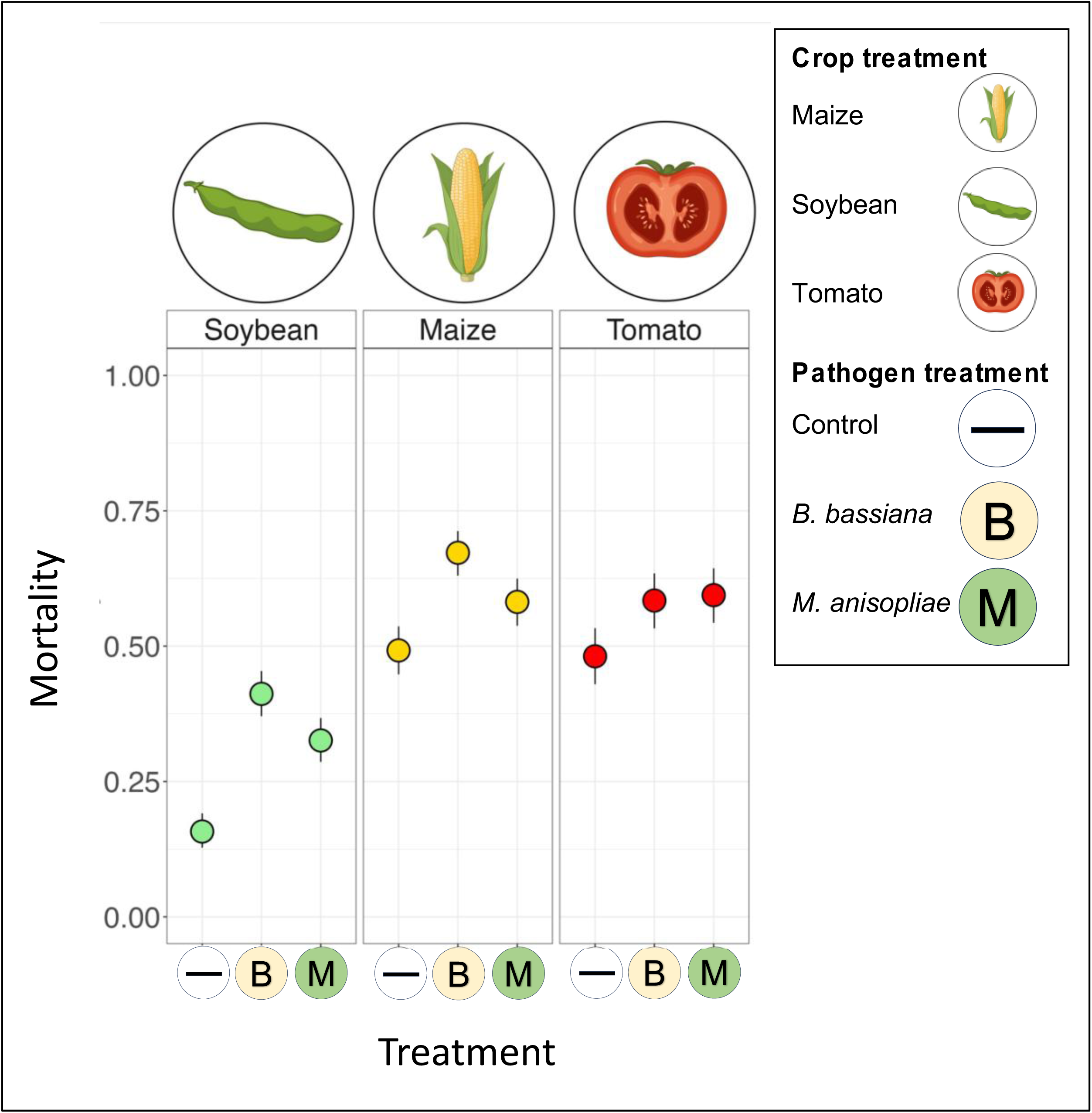
Relative ability of fungal isolates to kill *Helicoverpa* larvae depended on crop leaf diet. Dots indicate mean mortality at day 14 post-infection; whiskers give 95% binomial confidence limits for each combination of pathogen treatment (on the x-axis) and plant leaf diet (in panels). Total n = 3811 across the 9 treatment combinations.

### Heritabilities for larval ability to survive pathogen exposure

Risks of pest resistance evolution in response to fungal biopesticide application may be greatest if pest populations harbour pre-existing additive genetic variation for infection susceptibility on which selection can act. The half-sib *H. armigera* families in our experimental design varied greatly in ability to survive fungal pathogen exposure, and our calculations of heritabilities in each pathogen-diet treatment indicate substantial standing genetic variation for survival. Our Bayesian analyses provide posterior distributions of all model parameters, and the dense regions of these distributions have more statistical support (see Methods for more details).

Most of the evidence under the posterior heritability distributions supports relatively large values, consistent with substantial heritability in all treatments (0.10-0.75, see Fig. 2 and supplementary results). Counterintuitively, the heritabilities for survival in the fungal infected treatments were not noticeably higher than those in the control treatment (for any plant diet), which suggests that the standing genetic variation we observed is not solely related to pathogen defence. Instead, the genetic variation seems also to relate to general performance in our lab conditions (e.g., feeding on leaves of the three crops provided) even in the absence of pathogens.

**Fig. 2.**
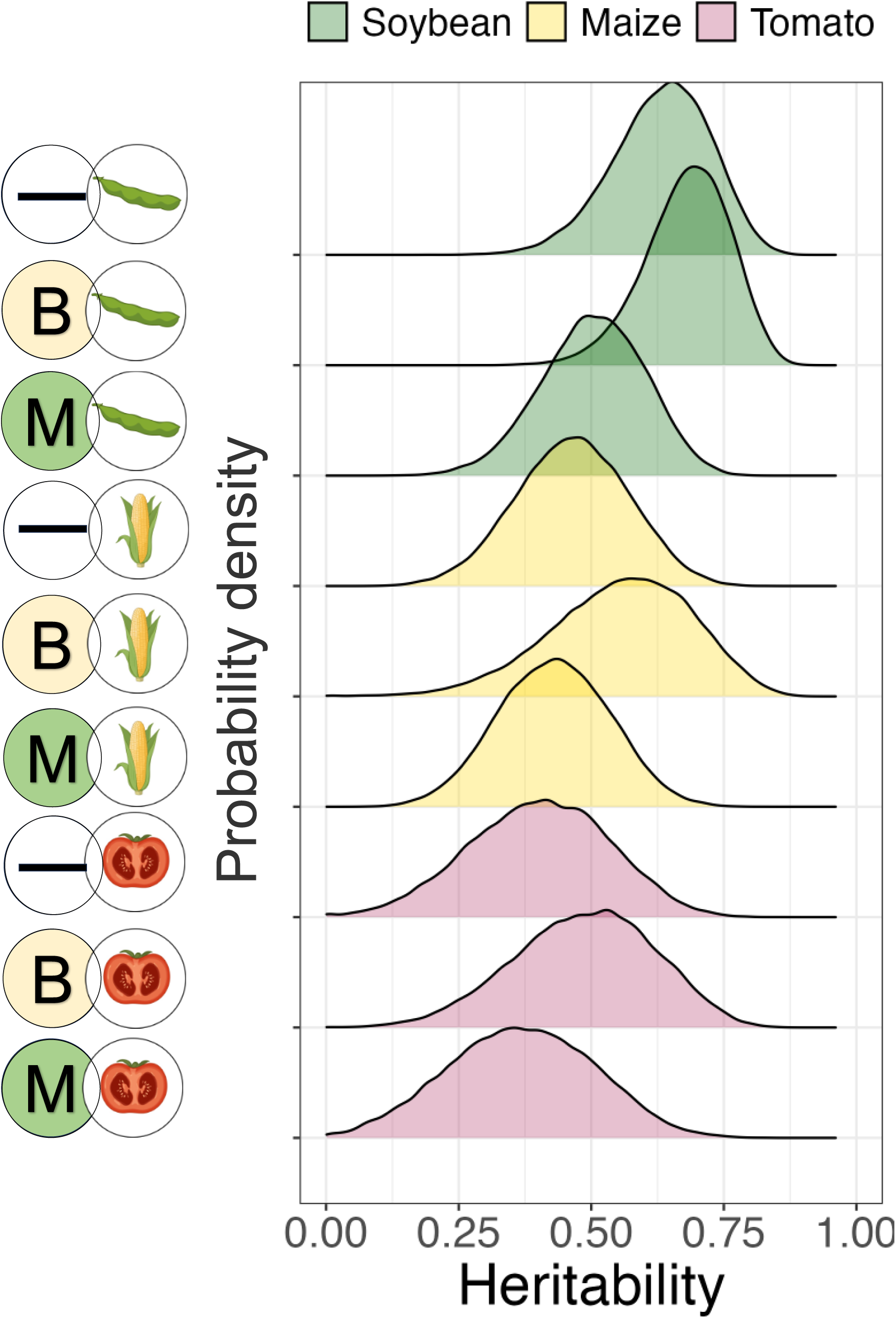
A substantial fraction of variation in larval survival in each treatment is heritable. Posterior distributions from a Bayesian analysis of heritabilities (the proportion of variation in survival that is additive and genetic) in each of the nine combinations of plant diet and infection treatment. The density of each distribution indicates the weight of evidence for a particular heritability value (on the x-axis). These estimates were obtained from 58 half-sib families of *H. armigera* fed on one of three leaf diets (soybean, maize, or tomato) and exposed to one of three pathogen-exposure treatments (M = *M. anisopliae*, B = *B. bassiana,* — = Control).

### Genotype by environment interactions for infection-survival

The high heritabilities we observed were driven by extreme survival differences between families: by 14 days post infection, in most pathogen treatments, families varied between some with close to 0% mortality and some with close to 100% mortality (Fig. 3). Those families that survived best under one combination of plant diet and infection treatment often performed relatively poorly in other treatment combinations. Indeed, the plant diet and pathogen treatments sharply affected the relative susceptibilities of families to infection, as expected if there are strong genotype-by-environment interactions affecting allelic fitness. We illustrate the change in family performance by ordering paternal families according to their mean survival whilst feeding on soybean and exposed to *B. bassiana*, plotted in the upper left panel of Fig. 3. By retaining the order of sire families across the remaining panels of Fig. 3, the neat cline in performance is only slightly disturbed by changes in pathogen treatment (left-most column of panels), whereas changes in plant diet (in the middle and right columns) have more disruptive effects on the rank performance of sire families. This visual inspection suggests that genetic correlations across treatments were lower (involving potential fitness trade-offs) when treatment contrasts involved changes in the diet than when they involve changes in pathogen treatment.

**Fig. 3.**
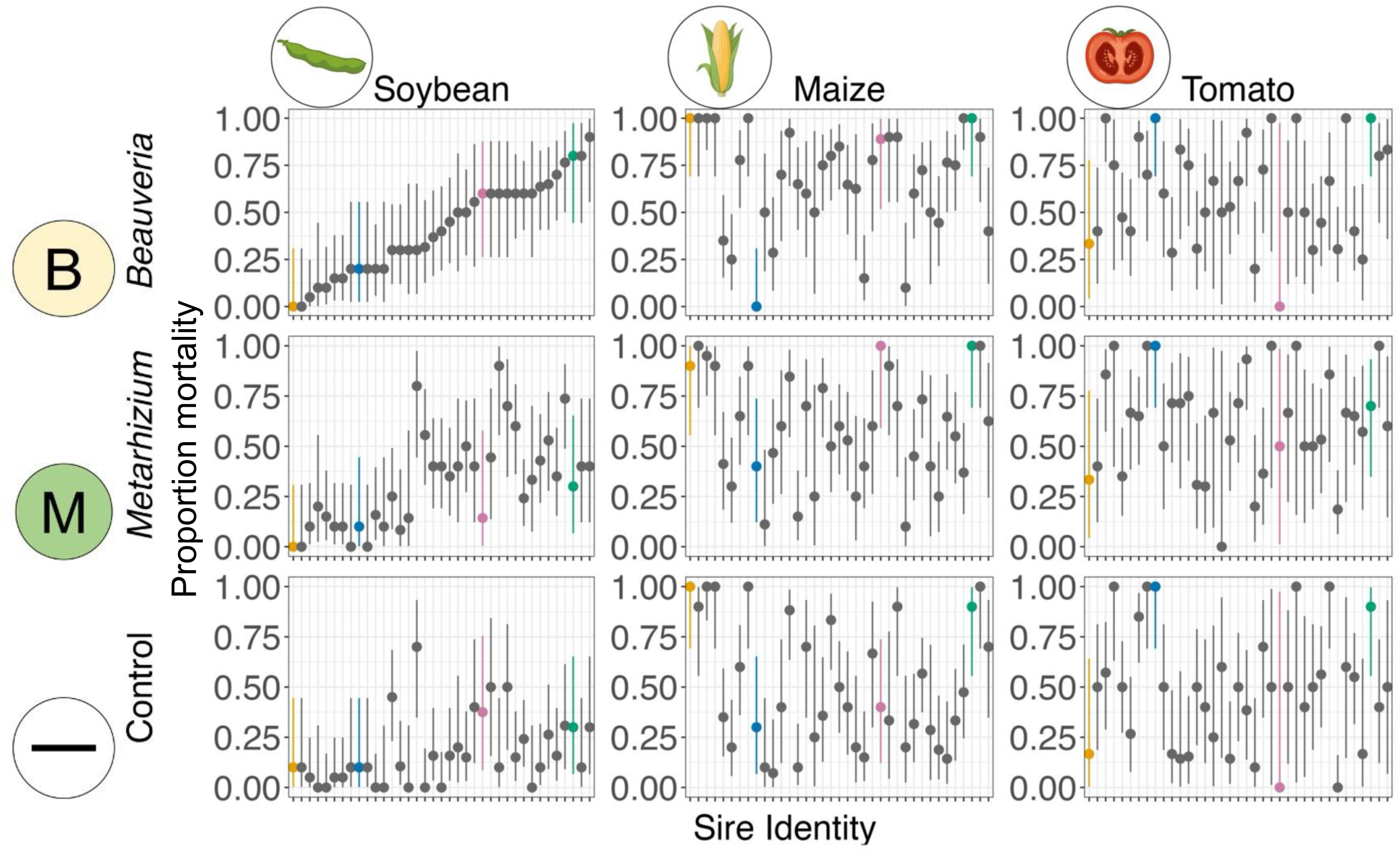
Diet and infection treatment strongly alter the relative fitness of different half-sibling families. Each point and whisker represents mean mortality and 95% binomial CI for sire half-sibling families 14 days post-exposure, depending on plant diet (columns) and pathogen treatment (rows). Four of these families are highlighted consistently across panels (see coloured points) to illustrate contrasting patterns of performance across habitats. The sires are arranged along the x-axis in order according to their rank performance when reared on soybean and exposed to *Beauveria*, as is clear in the cline in the upper-left panel (families at far left have the lowest mortalities, while those at far right have greater mortalities in the upper-left panel). The extent to which this cline is disturbed in different panels illustrates the extent to which family-level survival after pathogen exposure can be altered by the treatment in question. The phenotypic patterns in this plot are formally decomposed into the additive genetic components in Table 2 and Fig. 4.

**Table 2.**
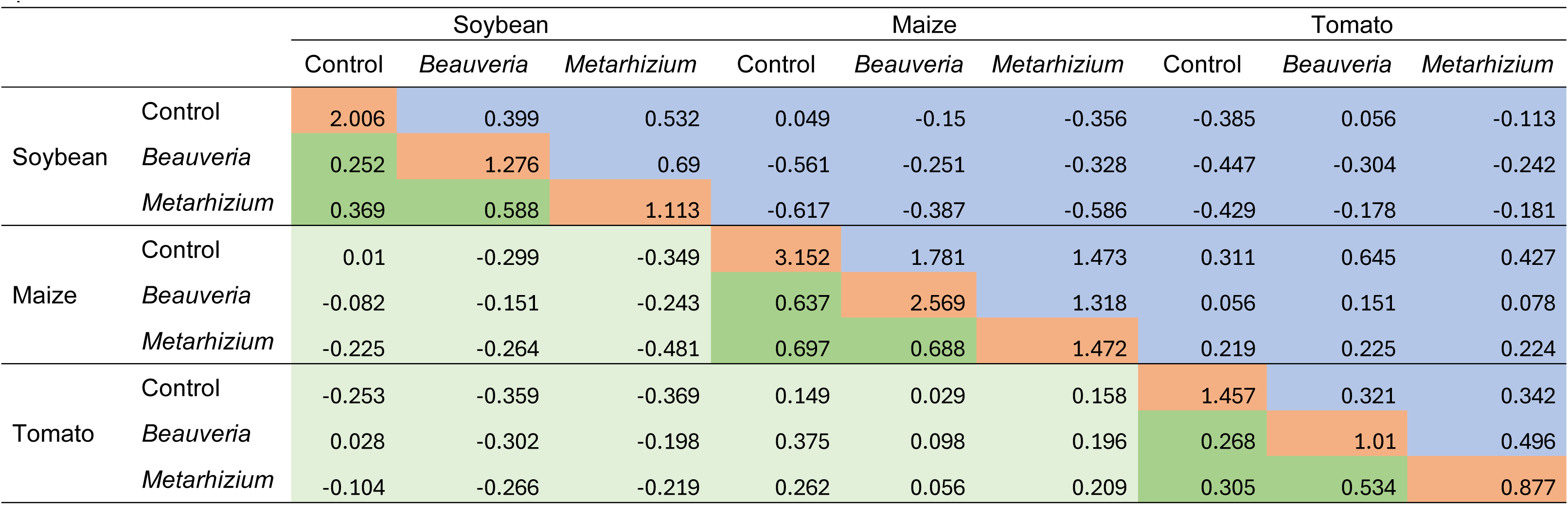
Changes in plant diet are associated with strong genetic trade-offs. Below are summaries of genetic variances (orange, diagonal), covariances (blue, upper off-diagonal), and correlations (green, lower off-diagonal) for mortality across 9 combinations of plant and pathogen exposure treatments. Within-plant diet correlations are shaded in darker green to call attention to their consistently higher values than the cross-plant diet correlations.

### Genetic correlations between larval survival ability in different crop-pathogen treatments

The results in Fig. 3 unexpectedly suggest that genotype fitness trade-offs driven by crop-diet might be stronger than those induced by changes in pathogen identity. To formally compare how crop and biopesticide changes affect the magnitude and direction of selection for biopesticide resistance, we calculated genetic correlations, which quantify the degree to which alleles for high fitness (larval survival to day 14) in one environment will also be selected for in a second. A perfect genetic correlation (*r*_g_ = 1) means that environmental changes do not alter relative allelic fitness. Correlations below one imply GEIs will slow responses to selection if environments change, and correlations below zero reverse the direction of selection^29^.

The magnitude and sign of cross-environment genetic correlations for survival depended markedly on whether the treatments differed in crop diet or pathogen exposure. Table 2 contains a summary of the single-model G-matrix (the matrix of variances and covariances) on the upper half-diagonal, and the genetic correlations on the lower half-diagonal. However, as for heritability estimates, the total evidence under the posterior is better illustrated by the ridgeplots in Fig. 4. Surprisingly, changing pathogen treatment without changing the host plant (either changing between fungal genera or from control to a pathogen infection treatment, Fig. 4, red shaded density ridges) depressed genetic correlations only modestly below 1 (mean = 0.48, bootstrapped 89% HDI = 0.39 -0.58). By contrast, changing the plant diet depressed genetic correlations far more, with many such correlations estimated below zero revealing crop-mediated genetic trade-offs for infection susceptibility (Fig. 4, green ridges, mean = -0.10, 89% HDI = -0.21 – 0.01). Counterintuitively, simultaneous change of both pathogen and diet treatments (right-most panel in Fig. 5) provided no obvious further depression in the genetic correlation (mean = -0.09, 89% HDI = - 0.17 – 0.01): when the genetic correlation involved environments that differ in both dimensions, the genetic correlations were barely distinguishable from those involving only plant diet changes.

**Fig. 4.**
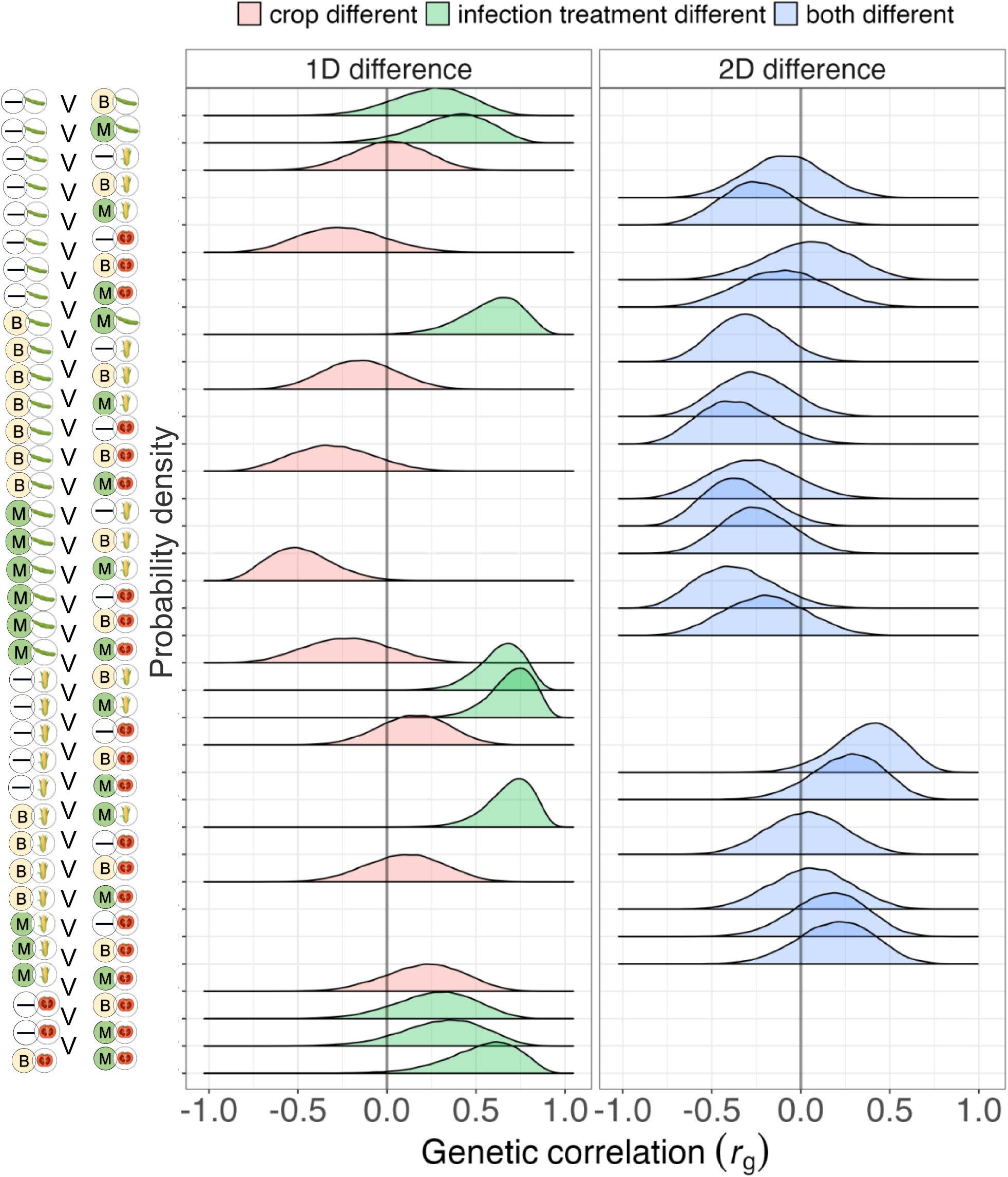
Genetic correlations are lower when measured across environments that differ in plant diet. Posterior distributions for cross-environment genetic correlations for mortality in *H. armigera* larvae grown in 9 different combinations of plant diet (soybean, maize, and tomato) and pathogen treatment (control, *Beauveria,* and *Metarhizium*). The 36 posteriors are clustered in the figure above depending on the axes of environmental difference, with environments differing in only 1 dimension on the left, and those differing in 2 dimensions on the right. Summaries of central tendency in the genetic correlations appear in the lower off-diagonal of Table 2.

**Fig. 5.**
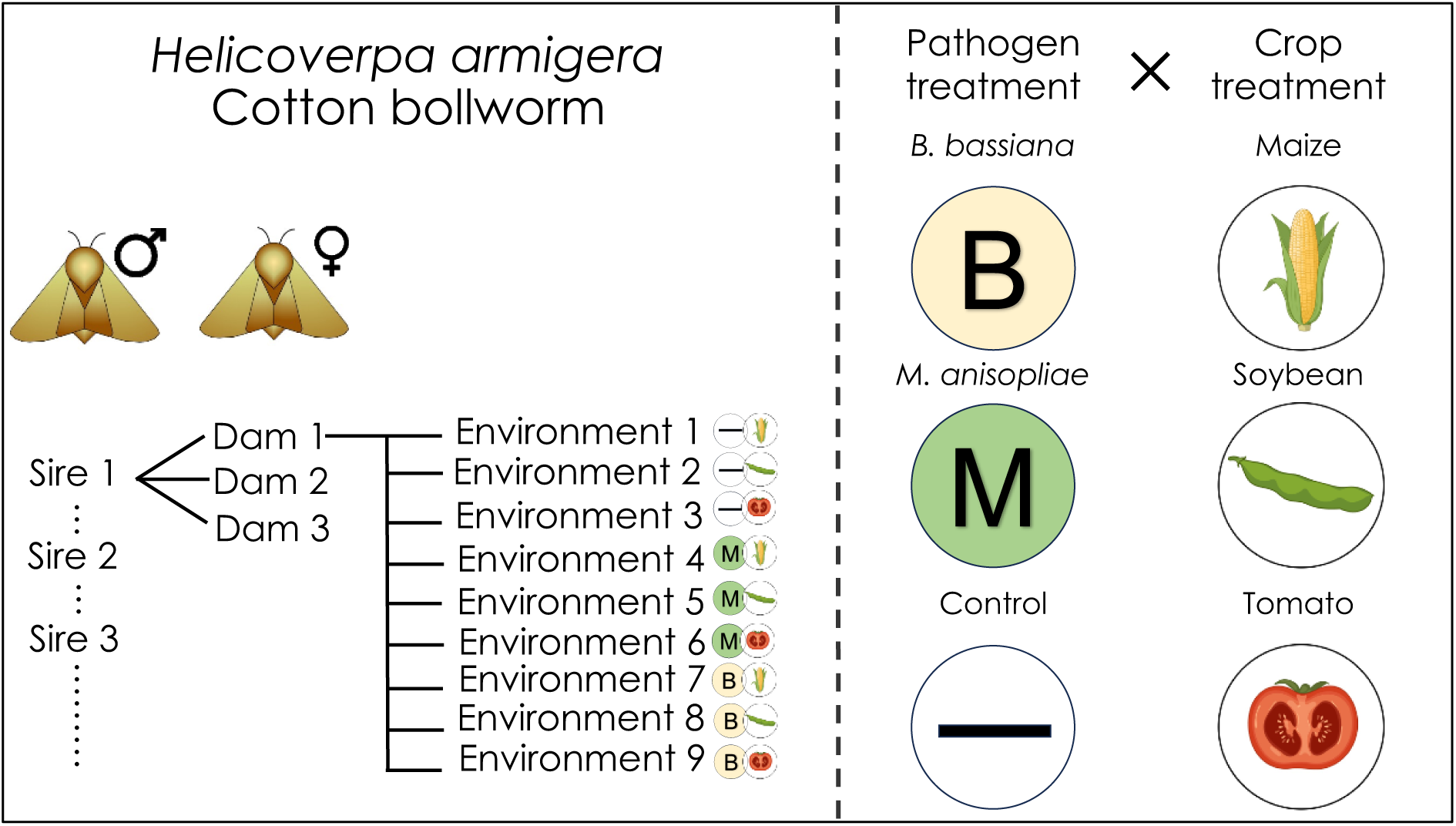
Schematic representation of the experimental design. Second instar larvae from each female were randomly assigned to one of 9 treatments. Insects were exposed to three different infection treatments and were reared on three different plants. Larval survival was recorded daily thereafter.

## Discussion

Pathogen exposure can exert strong selection on host populations. However, the extent to which this selection is shaped by external factors, such as host diet and pathogen identity, is not generally clear. Motivated by the need to assess how emerging risks of biopesticide resistance evolution could be managed, our study aimed to assess whether diet and pathogen differences drive GEIs for infection susceptibility. We found that genotypes of *H. armigera* vary substantially in their ability to survive infection by fungi that are used as biopesticides, as demonstrated by considerable heritabilities in all combinations of crop and pathogen treatment.

This high level of standing genetic variation presents a clear risk of resistance evolution against fungal biopesticides used in agriculture. We also reveal that altering the crop that larvae feed on can generate strongly inconsistent selection for resistance (evidenced by frequently negative genetic correlations for larval survival across plant diets), whilst changes in the identity of the fungal pathogen (where we found moderately positive genetic correlations) alter selection to a lesser extent.

The crop plant on which *H. armigera* larvae were feeding had a marked effect on survivorship. Larval survival in the uninfected control treatment was high on soybean leaves; however, approximately 50% of larvae died during two weeks on both the maize and tomato leaf diets. *H. armigera* feeds on over 100 plant species but not all support its survival, indeed secondary phytochemicals and nutrient deficiencies can impair fitness^30^. Some polyphagous herbivorous insect species, like *H. armigera,* are composed of many different genotypes that specialise on different plant species^31^, leading to genetically distinct populations based on crop type^32^. The population of *H. armigera* we studied may be better adapted to feed on soybean than the other two diets. We observed significant heritability for the ability to survive on all three plant diets under control conditions; this mirrors previous work studying *H. armigera* larval development on different chickpea varieties^33^. When we exposed larvae to *B. bassiana* or *M. anisopliae* fungal pathogens mortality rates increased compared to the control treatment. However, the magnitude of this infection-induced mortality was dependent on the precise combination of pathogen and plant diet: *B. bassiana* caused higher mortality than *M. anisopliae* when larvae fed on soybean and maize, whereas for larvae feeding on tomato leaves the two pathogens caused similar mortality. It is not clear why diet should modulate host infection susceptibility differentially depending on the identity of the pathogen; however, diet composition is well established to influence disease resistance^34^. Our data suggest that, for polyphagous pests, the efficacy of particular biopesticides may vary depending on the crop species a farmer grows, an observation that may have important consequences for biopesticide registration and use by the agricultural industry.

Pest insects frequently evolve resistance against chemical pesticides and genetically modified crops, yet the potential for resistance evolution against biopesticides formulated from living pathogens remains underappreciated, despite notable examples^7,35,36^. The risk of resistance will depend to a large extent on the existence and magnitude of pre-existing genetic variation for pathogen susceptibility, but such measures are unavailable for most pests. We demonstrate substantial heritabilities (the fraction of phenotypic variation due to genes) for pathogen susceptibility, but also show that these measures vary depending on the precise combination of pathogen treatment and plant diet. We note that the magnitude of heritability for survival in these pathogen exposure treatments is not solely driven by genetic variation for infection susceptibility, because we also observed substantial heritabilities in the absence of infection in the control treatments. Moreover, heritabilities in the wild, under heterogeneous field conditions will certainly be lower than in our standard laboratory conditions. Regardless, our results clearly support considerable standing genetic variation for survival in the presence of biological antagonists, and justify further work to quantify the risk of resistance evolution to biocontrol agents in this species and others under variable conditions^37^.

Classic concepts in host-pathogen evolution often assume that the ability to defend against one pathogen genotype comes at the cost of impaired ability to defend against others^38^. In the context of biopesticides, such trade-offs would mean that rotations of agricultural products containing different pathogens might be a highly effective resistance management approach^7,36^. Despite this *a priori* expectation, we were surprised to observe little evidence for pathogen-mediated susceptibility trade-offs. Our analysis revealed universally positive genetic correlations for host fitness between infection treatments containing either *B. bassiana* or *M. anisopliae*, meaning that insect genotypes best able to defend against one pathogen were generally well-equipped to defend against the other. We acknowledge that these microbes are both fungi, and that had we studied a wider range of pathogens of increasing phylogenetic distance our result may have been different. Nevertheless, our data indicate that host defence mechanisms against these fungal pathogens are generally broad, rather than specific to particular fungi. Some studies in *D. melanogaster* have similarly demonstrated positive correlations between defence against pairs of pathogens: *M. anisopliae* and *Pseudomonas aeruginosa*^20^, *B. bassiana and Lysinibacillus fusiformis*^39^. This lack of defence-specificity may stem from the absence of a closely coevolved relationship between host and parasite in these cases, a situation likely common among entomopathogenic fungal biopesticides used in crop pest control, as well as in the case of newly emerging infectious diseases.

Although there was no evidence of trade-offs between defence against *B. bassiana* and defence against *M. anisopliae*, the genetic correlations in susceptibility between these two pathogens were not very strong (i.e., very far from a perfect correlation of 1). The rank order of genotype susceptibility was only modestly conserved between the pathogens, which is clear evidence for pathogen-mediated GEIs. Thus, even in the absence of strong trade-offs, evolutionary responses in populations exposed to different fungal pathogens in sequence will be less rapid than when selection is driven consistently by a single pathogen genotype.

Evolutionary ecology theory frequently assumes that effective parasite defence is costly, and therefore not favoured by selection when pathogens are only encountered infrequently. Indeed, compelling evidence for this assumption exists for some parasites^40^. However, the mechanistic basis of resistance varies greatly across host-parasite systems, and not all mechanisms of resistance need be generally costly. If strong broad costs for infection defence occurred in the *H. armigera* – fungus system, these might be evident as negative genetic correlations between the control and fungus-exposed treatments. Surprisingly, the genetic correlations for survival are roughly the same whether they are between treatments that are two different pathogens, compared to cases where the contrast is between a pathogen treatment and the control treatment. From an applied perspective, this predicts that farmers rotating between biopesticides containing different microorganisms would be just as evolutionarily sustainable as alternating periods of biological control with periods of no pest control at all. If this finding proves to be general, there may be no benefit to biopesticide-free refugia on a sufficiently diverse landscape treated with multiple biopesticides. However, it is worth remembering that we modelled a single and simplified response variable: the ability to survive 14 days after infection. One might rightly question the extent to which this response adequately reflects selection across the entirety of larval development, and we invite more work on the genetic architecture of life history traits across multiple environments, even as we respect the appreciable samples and processing time that such experiments and analyses will require.

Host-parasite theory focuses on parasite identity, and to a lesser extent environmental temperature as drivers of inconsistent selection which prevent genetic variation for infection defence being efficiently purged by selection^3^. However, our data demonstrate that the importance of diet variation in generating heterogeneous selection for parasite resistance may have been greatly underappreciated. In striking contrast to the modestly weakened genetic correlations across pathogen treatments, genetic correlations for post-infection survival were strongly depressed by changes in plant diet and were far more likely to produce negative genetic correlations.

Indeed, shifting from soybean to another host plant consistently produced the most pronounced negative genetic correlations observed in our study. Genetic correlations for survival between maize and tomato diets were also low, but not as negative as those involving soybean and another crop. Whether this pattern is due to dietary specialisation for soybean in our moth population remains unclear, given the limited number of plant species we examined. Clearly, the specific nature of the habitats and pest populations will dictate whether trade-offs across heterogeneous patches can reverse biopesticide resistance evolution, and to what extent. In contrast to our findings, a study on aphids and their wasp parasitoids found little support that plant species altered the resistance of particular aphid genotypes to parasitism^41^. There are still too few studies of diet-induced GEIs to generalise, but the exciting possibility for inconsistent selection on parasite defence driven by diet variation suggested by our work invites further study.

What phenotypic differences account for observed survival variations among genotypes, and what biological process could explain why the fitness of genotypes to defend against infection is strongly dependent on the crop diet on which larvae feed? Microbial symbionts can have strong effects on the ability of insects to defend against infection^41^ and to feed on particular plant diets^42,43^. However, our estimates of genetic (co)variances come from the male parental contribution to offspring phenotypic variation; as we expect that most gut, or other, microbial symbionts would be maternally inherited^44^, we think that any non-genetic microbiome contributions to our estimates are probably small. Instead, a possible mechanism for diet-induced changes in pathogen resistance involves macronutrient-sensitive biochemical pathways. For example, dietary protein: carbohydrate ratios influence the ability of insects to upregulate immune responses and survive infection^34^. However, the extent to which these dietary effects vary between host genotypes to contribute to GEIs remains unclear.

In demonstrating that crop heterogeneity can alter the intensity and direction of multivariate selection for biopesticide resistance, we have answered a global food security question using theory from fundamental ecology and evolutionary science. Our research has important implications for agriculture and points to some outstanding questions requiring further research. We previously proposed that farmers could combat threats of biopesticide resistance evolution by engineering additional environmental heterogeneity into agricultural landscapes^7^; especially through use of spatial matrices or temporal rotations of different biopesticides and crops. In the present study, we show that some aspects of landscape heterogeneity could change the intensity and direction of selection on crop pests for defending against biopesticides but that not all dimensions are equally effective. For instance, our findings show modest negative genetic correlations across some plant diets, with an average of –0.10 for changes in plant diet alone, and a statistically indistinguishable –0.09 when diet differences are combined with different pathogen treatments. If biopesticide resistance were to start evolving in pest populations, the weakness of these negative genetic correlations indicates that reversing genetic adaptations through multivariate genomic evolution may require several generations in varied habitats. This underscores the need for a broad spectrum of divergent selection strategies to maintain effective and sustainable pest control, beyond the limited number of pest control products that were initially envisioned to induce negatively correlated cross resistance for chemical pesticides^18^.

Whether the many dimensions needed to manage the risks of resistance evolution to biocontrol already exist in some complex agricultural landscapes, or whether they need to be augmented through active diversifying efforts, and at what scale, is not yet clear. Undoubtedly, the answer will depend on the extent to which pests disperse, the genetic architecture for surviving in distinct habitat patches, pest demographics, as well as local landscape management practices. Many of the most destructive crop pests are invasive precisely because they can disperse long distances, and in those species only coarse spatial heterogeneity may be needed to sustain genetic variation. For pests that disperse much less, temporal heterogeneity may be more important to help prevent local adaptation. In landscapes dominated by individual small-holders (routinely exhibiting landscape diversity at relatively small scales), highly dispersing pest populations will experience gene flow across farms, so there would be no need for individual farmers to create additional on-farm diversity merely to reduce directional evolution (although there could be ecological benefits for doing so). In contrast, in landscapes dominated by extensive industrial monocultures, enhancing spatial and temporal diversity, by favouring shifting mosaics^7^, perhaps involving coordination among farmers, will likely be more important.

This research illustrates how the principles of genetic variation and GEIs, fundamental to evolutionary ecology are crucial not only for understanding natural systems but also for devising effective, sustainable management strategies in agriculture. Such insights underscore the importance of integrating ecological and evolutionary science to tackle complex global challenges like food security, highlighting the need for a deeper exploration of the interactions between pest genetics, landscape structure, and management practices. In addition, our work significantly advances understanding of ecological and evolutionary dynamics in pest populations. Our results bolster the theoretical prediction that divergent selection is a pivotal force in maintaining genetic variation for key life history traits. Importantly, our work calls into question the prevailing expectation that trade-offs in host resistance to different pathogens are primarily responsible for maintaining genetic variation for pathogen defence traits. Instead, we show that other biological factors, including plant diet (or prey more generally, perhaps), have a mediating role in selection for pathogen resistance. Because organisms occupy complex and heterogeneous habitats, in which multiple biological antagonists exert temporally and spatially variable selection on many alleles, changes in the broad biological environment are probably more important for the maintenance of diversity in all life history traits, including infection defence, than is generally acknowledged in the host-pathogen literature.

## Methods

### Plants

*Soybean* (*Glycine max* (variety Summer Shell)), tomato (*Solanum lycopersicum* (variety Roma)) and maize (*Zea mays* (variety Tramunt)) were grown from seed (Tamar Organics, UK) in a controlled environment facility at the University of Stirling (16:8 hr L:D photoperiod with compact-fluorescent lamps; 24°C/16°C during L/D; 70% R.H.). Seeds were placed individually in small pots (5cm × 5cm × 5cm) containing approximately 150 g John Innes Seed Compost to germinate. Germinated seeds were then transferred to larger pots (12cm × 15cm) in approximately 700g John Innes No 2 compost.

### Preparation of fungal material

We used two fungal isolates from Campinas Biological Institute (Brazil) that are virulent against *H. armigera* (IBCB 1363 (*B. bassiana*) and IBCB 425 (*M. anisopliae*)^45^. Fungal material was grown on potato dextrose agar with chloramphenicol (5 × 10^−5^ g ml^−1^). Agar plates were incubated for 10 days (25°C, 24 hr dark), then dried at room temperature for approximately 10 days; plates were rotated periodically to ensure even drying. Then, sporulating fungal material was scraped from the plates and spores dried further on silica gel in a fridge before being suspended in sunflower oil. These formulations were vortexed, and then briefly agitated with a probe sonicator to break up spore masses. Spore suspension concentrations were determined using a haemocytometer and adjusted to 2 × 10^7^ conidia ml^-1^.

### Experimental system

All insect culturing and experiments were conducted in the quarantine facility at the University of Stirling in controlled environment rooms. *Helicoverpa armigera* pupae were sourced from Andermatt Biocontrol AG, Switzerland. The insects used in this experiment originated from six separate consignments of pupae from Switzerland. On arrival, pupae were washed in 1% (w/v) copper sulphate solution, sexed and placed under conditions of reversed photoperiod (10 hrs dark between 03.00 and 13.00 hrs). Male pupae were held at 27°C, and females at 25°C to hasten the emergence of adult males and ensure sexual maturity synchrony of the sexes.

Single mating pairs (one female <24HRS old and one male >3 days old) were placed in ventilated plastic containers (55 mm (l) × 55 mm (w) × 60 mm (h)) and provided with vitamin solution^46^. Males always originated from the shipment preceding the females to ensure outcrossing.

### Experimental design

The experimental design (Fig. 5) involved studying survival of half-sibling larvae in each of nine different experimental treatments (combining three diets, two pathogen infection treatments and an uninfected control). To examine genetic variation in defence traits, we mated each of 37 sires with up to three dams, resulting in 37 paternal half-sib families^47^.

Mating pairs were observed for female receptivity to males: for females, calling (pheromone release) signified the attainment of reproductive maturity, a behaviour which was immediately identifiable and characterised by the female’s extruded ovipositor. The maturity status of males was tested by examining their response to a calling female. Male mating behaviour consisted of brush extension and swiping movements of the abdomen directed at the calling female. Males that did not attempt to mate were classified as immature and tested again in 24 hrs.

Unreceptive females moved away from an approaching male, withdrew the ovipositor, and flexed the abdomen resulting in the tip held beneath the female and inaccessible to male claspers. If a male was successful grasping an unreceptive female’s abdomen, females initiated violent wing fanning to escape. This behaviour starkly contrasted with that of receptive females, who ceased all activity about 15 seconds after pairing with a male. Mating pairs were allotted 15 min to mate. If unsuccessful, moths were repaired and observed as before. Mating was deemed successful if the moths remained attached for more than 20 min and then separated successfully. Females that had successfully mated were transferred to ventilated plastic boxes (55 mm (l) × 55 mm (w) × 60 mm (h)) for oviposition and fed on cotton wool soaked with vitamin solution^46^.

*H. armigera* egg masses were collected daily from each female, split approximately evenly into three groups, and placed in ventilated plastic boxes (7cm × 7cm × 7 cm) containing either fresh maize, soya or tomato leaves until hatching. Fresh leaf material was added daily. Early second instar *H. armigera* larvae from each plant treatment were randomly assigned to one of three infection treatments with sunflower oil spore suspensions: *B. bassiana* IBCB1363, *M. anisopliae* IBCB 425 and a pathogen-free control. Larvae were placed individually in Petri dishes (4.5 cm diameter) and 0.5µl spore suspension (or blank oil) was pipetted onto the larval cuticle. Larvae were left for 2 hrs, then transferred to individual *Drosophila* vials (23ml, Sarstedt), sealed with breathable cellulose acetate flugs and provided with fresh leaves of the plant treatment they had previously fed on. Larvae were held at 25°C, 75% R.H. and 14:10 hr L:D). Mortality of larvae was first recorded 24 hrs after treatment and then daily thereafter until death or pupation; larvae were transferred using a sterilized fine brush onto a fresh leaf diet when required. This experiment was conducted over two blocks, with identical experimental design for each; block 1 used 18 sires (32 dams), and block 2 had 19 sires (26 dams).

### Data Analysis

From each dam, we attempted to rear 90 offspring, though in some cases we obtained fewer than the necessary number of eggs. Consequently, our preliminary offspring count was 4344 larvae, of which 4314 survived to day 3 when the infection/control treatment was applied. To avoid error in quantifying mortality variation that was not due to pathogens, we excluded a small proportion of larvae that died shortly after inoculation in a way that was unlikely to reflect pathogen infection. For example, in some cases (7.8%, N = 339) death occurred within two days of treatment, a time when fungal infection is unlikely to have proceeded to the lethal stage. In other cases (3.7%, N = 162), the larvae never moved after being treated, and so may have died immediately following oil application. To prevent these instances from obscuring patterns that were due to the treatments, we removed both categories before analysis. A small number of larvae (N = 2) also escaped before treatment. This left us with 3811 larvae included in the final dataset.

We performed all statistical analyses using R4.3.2^48^. We computed mortality rates at daily intervals from day 5 after infection through day 14 (a range that should capture most of the relevant pathogen-induced mortality) and we chose the day on which distinctions between control treatments and pathogen treatments were highest (day 14), to most closely capture differences in genetic variation for pathogen susceptibility.

To describe patterns of mortality in pathogen and plant diet treatments, we built a generalised linear model with logit link implemented in lme4^49^ in which both pathogen and plant-diet treatments as well as their interaction were predictors of the binomial survival proportion. We fitted random effects for both sire and dam identity to account for non-independence of larvae due to family membership and to control for maternal effects. We used parametric bootstrapping^50^ to test the significance of the plant:pathogen treatment interaction when comparing nested models.

To compute quantitative genetic parameters, we fit generalised models using the brms package^51^. We fitted the combination of plant diet and pathogen treatment as a 9-level fixed factor and fitted insect sires and dams as random effects. We further allowed the effect of sire to vary by treatment and asked the model to estimate correlations across treatments to reconstruct the G-matrix. We did not similarly allow dam effects to vary by treatment, because effects related to maternal condition (e.g., through egg provisioning) should uniformly improve offspring survival regardless of the specific treatment. Note that in our experiments dams were never exposed to pathogens and were fed an artificial diet, so there is no potential for interesting trans-generational maternal advantages due to phenotype matching. To ensure sufficient warmup and chain mixing, we ran the models for 32,000 iterations (half of which were used as warmup iterations) and adjusted the NUTS sampler by setting adapt_delta to 0.96 to avoid divergent transitions. Our models produced well-mixed chains and unimodal posterior distributions. The estimates were robust to minor changes in prior specifications, and were not influenced noticeably by alternate model structures (e.g., fitting only two treatments to produce a single genetic correlation estimate instead of estimating all 36 estimates simultaneously from a single model).

We used samples from the posterior to compute heritabilities within each combination of plant diet and pathogen treatment. For each sample, the environment-specific heritability was calculated as the sire variance in that environment divided by the sum of the sire variance, the maternal variance, and the residual variance (fixed at 1 because this is a generalised model) in that posterior draw. We also used posterior samples to compute genetic correlations; these are extracted for each pair of environments, and since there are 9 environments there are 36 pairwise combinations. We analysed patterns for the genetic correlations with respect to change in plant diet, pathogen treatment, and the combination of the two. To highlight the Bayesian nature of these analyses we report 89% highest posterior density intervals (89% HPDI) when comparing different environmental contrasts, but we note that the complete posterior distribution is the best representation of *a posteriori* evidence^52^. For this reason, to facilitate an appreciation of the total evidence we illustrate posterior densities using ridgeplots^53^.

## Acknowledgements

All authors were supported by a joint Newton Fund international partnership between the Biotechnology and Biological Sciences Research Council (BBSRC) in the UK and the São Paulo Research Foundation (FAPESP) in Brazil under BBSRC awards reference BB/R022674/1 & BB/S018956/1 and Grant 2018/21089-3, São Paulo Research Foundation (FAPESP). Additionally, LFB was supported by grants from Vetenskapsrådet (Sweden): 2021-05466, and the Carl Trygger Foundation (20:63). We are grateful to James Weir for technical support and management of controlled environment facilities.

## Author contributions

Conceptualization: MCT, RAP, LFB

Methodology: RM, MCT, EF, LFB

Formal Analysis: RM, MCT, LFB

Investigation: RM, EF

Writing—original draft: RM, MCT, LFB

Writing—review & editing: RM, MCT, RAP, LFB

Visualization: RM, MCT, LFB

Supervision: MCT, LFB

Project Administration: RM, MCT, LFB

Funding Acquisition: RM, MCT, RAP, LFB

## Data Availability Statement

Data and code will be uploaded to a freely accessible online repository on acceptance.

## Extended Data

**Table S1.** Parameter estimates from a generalised linear mixed model (with logit link) of larval survival as a function of food plant and fungal isolate treatment. The interaction p-value was estimated using parametric bootstrapping to compare models with and without the interaction term. Note that the residual variance for a binomial model is fixed at 1.

| term | estimate | Std. error | z-statistic | pb P-value |
| --- | --- | --- | --- | --- |
| (Intercept) | -1.802 | 0.162 | -11.137 |  |
| Crop (Maize) | 1.802 | 0.155 | 11.661 |  |
| Crop (Tomato) | 1.856 | 0.166 | 11.165 |  |
| Isolate (*Beauveria*) | 1.423 | 0.152 | 9.353 |  |
| Isolate (*Metarhizium*) | 1.004 | 0.155 | 6.462 |  |
| Crop (Maize): Isolate (*Beauveria*) | -0.607 | 0.203 | -2.991 | 0.003 |
| Crop (Tomato): Isolate (*Beauveria*) | -0.962 | 0.217 | -4.438 |  |
| Crop (Maize): Isolate (*Metarhizium*) | -0.621 | 0.204 | -3.043 |  |
| Crop (Tomato): Isolate (*Metarhizium*) | -0.476 | 0.219 | -2.171 |  |
| Maternal random sd | 0.517 |  |  |  |
| Paternal random sd | 0.463 |  |  |  |

